# Caloric restriction overcomes pre-diabetic and hypertension induced by high fat diet and renal artery stenosis

**DOI:** 10.1101/2021.03.30.437684

**Authors:** Michelle Sabino de Souza Nunes Faria, Vinicíus Eduardo Pimentel, Júlia Venturini Helaehil, Mayara Correa Bertolo, Nathalia Tonus Horas Santos, Pedro Vieira da Silva-Neto, Bruna Fontana Thomazini, Camila Andréa de Oliveira, Maria Esméria Corezola do Amaral

## Abstract

**Background:** Caloric restriction (CR) is a type of dietary intervention enjoyed as an essential tool in weight loss by modulating critical pathways of metabolic control, although it is not yet clear what repercussions this intervention model results when associated with renovascular hypertension. Here we demonstrate that CR can be beneficial in obese and hypertensive animals.

**Methods:** Rats were divided into groups: SHAM, and two groups underwent surgery to clip the left renal artery, to induce renovascular hypertension (OH and OHR). The SHAM diet was performed: 14 weeks normolipidic diet; OH: 2 weeks normolipidic diet + 12 weeks hyperlipidic diet, both ad libitum; OHR: 2 weeks normolipidic diet + 8 weeks ad libitum high fat diet + 4 weeks restricted 40% high fat diet.

**Results:** the OHR group dissipated blood pressure, body weight and glucose homeostasis. Reductions in insulinemia, lipids, islets fibrotic areas in the OHR group were observed along with increased insulin sensitivity and normalization of the insulin-degrading enzyme. Nicotinamide phosphoribosyltransferase, insulin receptor, Sirtuin 1 and complex II protein were modulated in liver tissue in the OHR group. Strong correlations, direct or indirect, were evaluated by Spearman’s model between SIRT1, AMPK, NAMPT, PGC-1α and NNMT with the reestablishment of blood pressure, weight loss, glycidic and lipid panel and mitochondrial adaptation.

**Conclusion:** CR provided short-term beneficial effects to recover physiological parameters induced by a high-fat diet and renal artery stenosis in obese and hypertensive animals. These benefits, even in the short term, can bring physiological benefits in the long run.

## Introduction

Obesity and hypertension have become important global public health problems, resulting in morbidity and mortality. Together, they are responsible for a 5-fold increase in the possibility of developing type 2 diabetes and secondary cardiovascular diseases [30]. The main causes of obesity and hypertension are related to food, which over the years has changed with the increase in the content of fats and carbohydrates. This, together with the lack of physical activity, provides conditions conducive to the appearance of metabolic disorders such as diabetes [7].

The pancreas and liver are essential for establishing homeostasis of carbohydrates and lipids. Insulin is secreted by pancreatic β cells in response to increased blood glucose levels after a meal. Insulin increases the rate of glucose uptake, especially in skeletal muscle and adipose tissue. The increase in insulin stimulates glycogen synthesis in hepatocytes through a reduction in GSK-3, which increases glycogen synthase, providing hepatic glycogen synthesis. The excess passes through the pathways that lead to the synthesis of triglycerides for transfer to adipose tissue. Insulin also reduces hepatic gluconeogenesis and hepatic glycogenolysis.

Therefore, a possible way of treating metabolic disorders is caloric restriction (CR), involving the use of a diet with a 25-60% reduction in nutrients, in relation to the free diet, with a low intake of calories derived from carbohydrates, fats and/or proteins, without causing malnutrition [17]. It is proposed that the CR acts to reduce oxidative stress, decreasing oxygen consumption and stimulating the intracellular signaling pathways that regulate the metabolism of glucose, lipids and proteins [16] found that CR regulated the expression of SIRT1 (regulator of the silent form of transcription 1), a molecule involved in the regulation of various pathological processes related to oxidative stress, anti-aging, cell survival and stimulates the formation of autophagosomes, increasing autophagy in conditions of food restriction [33]. There is also evidence that SIRT1 acts in the regulation of glucose and lipid metabolism [19], however there is still no evidence to demonstrate the antihypertensive effects of CR concomitantly with the activity of SIRT1.

SIRT1 requires nicotinamide adenine dinucleotide (NAD+) for its enzymatic activity. Studies have shown that Nampt (nicotinamide phosphoribosyltransferase) is a limiting enzyme in the biosynthetic pathway of NAD+ in mammals and directly regulates the activity of SIRT1. Nampt and NAD+ levels in tissues and organs are reduced by a diet high in fat and aging, contributing to type 2 diabetes [18] levels of transcription and expression of Nampt were down-regulated in rodent hearts in response to cardiac hypertrophy induced by pressure overload, while overexpression of Nampt in cardiomyocytes increased cellular levels of NAD+ and ATP [27].

The regulator of the silent form of transcription 3 (SIRT3), predominantly mitochondrial is also involved in CR, as it increases the demand for NAD+ in the mitochondria through Nampt, protecting against oxidative stress and cell death and suppressing the generation of reactive oxygen species [26]. Protection is provided against cellular aging by regulating glucose metabolism [21]. Taken together, these data also suggest that CR promotes, in addition to glycidic homeostasis, an anti-aging action guided by increased autophagy and reduced oxidative stress, establishing itself as a beneficial experimental model.

In the central physiological response to the action of CR, the liver and pancreas play a leading role, as they are central organs for alterations in the metabolism of lipids and glucose in response to nutritional and hormonal signals. Here, we focus our studies on a better understanding of how these organs act in conditions of caloric restriction and renovascular hypertension in obesity in animals, in order to elucidate the role of this dietary intervention in two pathologies that negatively interfere in glycemic and lipid homeostasis.

## Materials and methods

### Chemicals and reagents

Bovine serum albumin (BSA, fraction V), tris-(hydroxymethyl)aminomethane (Tris), phenylmethylsulfonylfluoride (PMSF), dithiothreitol (DTT), Triton X-100, Tween 20, aprotinin, and glycerol were purchased from Sigma-Aldrich® (St.Louis, Missouri, USA). The reagents and apparatus for sodium dodecyl sulfate-polyacrylamide gel electrophoresis (SDS-PAGE) and immunoblotting were from Bio-Rad® (Richmond, California, USA). PVDF membranes (BioRad®) and the SuperSignal West Pico Chemiluminescent Substrate Western blotting analysis system were from Thermo Fisher Scientific Inc®. (Rockford, Illinois, USA). Glucose was purchased from Fluka Biochemika® (Buchs, Switzerland) and NPH human insulin (Novolin® N – Novo Nordisk, Bagsvaerd, Dinamarca).

### Animals

Male *Wistar* rats (50-days-old) were used for in vivo experiments. The rats were paired by weight and age for the start of the experimental protocol and in all experiments (n=5 each group). The animals from the Vivarium of the University Center of the Hermínio Ometto Foundation, where maintenance and experiments were carried out in accordance with the institutional guidelines for ethics in animal experiments approved by the Center for Animal Research Ethics and Research (CEUA) of the Hermínio Ometto de Araras Foundation (protocol number 042/2016).

The animals were divided into three groups: SHAM, OH (hypertensive and obese), and OHR (hypertensive, obese, and with 40% caloric restriction). Renovascular arterial hypertension was induced by clipping of the left renal artery, using the Goldblatt 2K1C technique [12].

### Diets and interventions

Part of the animals had access to the standard diet with commercial Nuvilab® food for rodents and others had access to the Open Source Diets® high fat diet. The animals in the SHAM group received, throughout the experiment, exclusively Nuvilab® normolipidic food *ad libitum*, while the OH and OHR group received this food only for 2 weeks *ad libitum*. After this period, and the matching of body weight between the animals of the different groups was verified, we started the obesity induction protocol by means of an Open Source Diets® hyperlipidic diet, for 2 months’ *ad libitum*. After this period, and the body weight matching was verified, the OH group continued to receive a high-fat diet for one-month *ad libitum*, while the OHR group, in this same period, had a 40% restriction of the group’s average consumption of OH. The composition of the diets is shown in Table 1.

**Table 1.** Nutritional composition of the diets.

|  | Hyperlipidic diet <sup>a</sup> | Normolipidic diet <sup>b</sup> |
| --- | --- | --- |
| Protein | 23.65% | 22.23% |
| Carbohydrate | 40.27% | 60% |
| Fiber | 5.80% | 5.73% |
| Fat | 23.60% | 4.0% |
| Minerals | 5.24% | 9.0% |
| Vitamins/supplements | 1.40% | * |
| Red dye | 0.006% | * |
| Energy | 4.72 kcal/g | 3.97 kcal/g |
| Calcium | * | 0.92% |
| Phosphorus | * | 0.87% |

Note: \* value not determined.
<sup>a</sup> Source: Research Diets Inc.
<sup>b</sup> Source: Nuvilab CR1-Nuvital<sup>®</sup>.

### Body and blood pressure parameters

The animals were weighed on a conventional digital scale, properly calibrated, throughout the experimental period.

The blood pressure was analyzed for plethysmography of tail. For this, a sleeve was coupled to a pressure transducer was put around the tail of animal awake, previously the animals were placed in a heating booth 37°C. Pressure variants was captured in specific data software: PowerLab a/S anolog-to-digital converter (AD Instruments Ltd., Csdtle Hill, Australia). The blood pressure was measure weekly during the experiment.

The Lee index was obtained as the ratio between the cubic root of the body weight and the nasoanal length of the animal (Buyukdere et al. 2019), described by the formula:

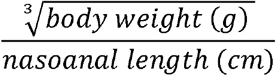

### Ip.GTT (intraperitoneal glucose tolerance test) and Ip.ITT (intraperitoneal insulin tolerance test)

The ip.GTT tests were performed after fasting the animals for 6h (n=5 animals per group). The animals were administered glucose (2g/kg body weight) intraperitoneally. Blood samples were collected from the tail, before glucose overload (time zero) and at 30, 60, 90, and 120 min after glucose infusion.

The blood glucose was determined using reagent strips and a glucose meter (Abbott®, Chicago, USA). The ip.ITT tests were performed after fasting the rats for 6h. The animals were administered standard crystalline insulin (1.5U/kg body weight), intraperitoneally. Blood was collected at time zero (before insulin injection) and at times of 5, 10, 15, 20, 25, and 30 min.

Blood glucose was determined using reagent strips and the Abbott® glucose meter. The glucose disappearance constant (Kitt) was calculated using the formula: ln2/t1/2. The serum glucose t1/2 value was calculated from the slope of the minimum regression curve, considering the linear phase of decrease of the plasma glucose concentration. The blood glucose was determined using reagent strips and the glucose meter.

### Determination of Hepatic and Muscle Glycogen Content

The gastrocnemic muscle and the liver of the animals were dissected and the determination of the hepatic and muscular glycogen concentration was done. Approximately 100mg of tissue was placed in 30% KOH and incubated at 100 °C for 15 min. Afterwards, 100% ethanol was added and incubated for 15 minutes at 70 °C. The test tubes were placed on ice for 20 min and then centrifuged for 10 min. The pellet was suspended in water and the glycogen was evaluated using a colorimetric method [20].

### Euthanasia and samples

After the trial, the animals were euthanized by deepening anesthesia with xylazine (10mg/kg) and ketamine (90mg/kg). Serum was collected for biochemical analyses. The liver and pancreas were removed for stereological, morphological, and biochemical analyses, as well as for protein quantification by Western Blotting.

The animals’ blood was collected during euthanasia via cardiac puncture, after an 8-hour fast. This material was centrifuged for 10 minutes at 3500 rpm to obtain the serum. After centrifugation, the serum was frozen until the time of analysis.

### Biochemical dosages

Serum levels of Aspartate aminotransferase and Alanine aminotransferase, total cholesterol and fractions, triglycerides, total protein and glucose with Biotécnica® commercial colorimetric kit, following the manufacturer’s instructions (Varginha, MG, Brazil). Insulin was determined by ELISA (Biotec Center, SPI-Bio and Ellipse Pharmaceuticals, France).

### Determination of cholesterol and hepatic triglycerides

Approximately 100mg of liver tissue was added to a 2:1 chloroform-methanol solution, crushed and stirred for 1 minute. Afterwards, methanol was added to the crushed and centrifuged for 10 minutes at 3000 rpm. Sequentially, the supernatant was transferred to a conical bottom tube containing chloroform and a 0.73% NaCl solution. Again centrifuged, the supernatant discarded, and the precipitate washed three times with Folch’s solution (3% chloroform, 48% methanol, 47% water and 2% 0.2% NaCl). The lipid extracts were dried in a ventilated oven at 37°C, resuspended with 1 ml of isopropyl alcohol for later determination of cholesterol and triglycerides using the commercial colorimetric kit Labtest®, following the manufacturer’s instructions (Lagoa Santa, MG, Brazil) [9].

### Western Blotting

The liver tissue was homogenized in protein extraction buffer, with aliquots being used for protein quantification according to the Biuret method. The samples were treated with Laemmli buffer (0.1% bromophenol blue, 1M sodium phosphate, 50% glycerol, and 10% SDS).

The electrophoresis runs were performed with 50μg of protein. The proteins were transferred from the gel to PVDF membranes (Immun-Blot, Bio-Rad®). After transfer, the membranes were blocked with basal solution containing bovine albumin and were then incubated with the primary antibodies (Table 2) overnight, at 4 °C, under agitation.

**Table 2.** List of the antibodies used.

|  |  |
| --- | --- |
| IR | Insulin Rβ (CT-3): sc-57342 (1:200) - Santa Cruz<sup>®</sup> |
| pAkt | Phospho-Akt (Ser 473): #9271 (1:1000) - Cell Signaling<sup>®</sup> |
| Akt | Akt Antibody #9272 (1:1000) - Cell Signaling<sup>®</sup> |
| Sirt1 | Sirt1 (C14H4) - 1:1000 - Cell Signaling<sup>®</sup> |
| Nampt | PBEF (E-3) sc-393444 - 1:200 - Santa Cruz<sup>®</sup> |
| Sirt3 | (C73E3) rabbit mAb - 1:200 - Santa Cruz<sup>®</sup> |
| NNMT | (G-4): sc-376048 - 1:200 - Santa Cruz<sup>®</sup> |
| pAMPKα | (Thr 172) - 1:1000 - Cell Signaling<sup>®</sup> |
| AMPKα | 2532# 1:1000 - Cell Signaling<sup>®</sup> |
| PGC1α | (4A8): sc-517380 - 1:200 - Santa Cruz<sup>®</sup> |
| Foxo | (D-12): sc-48348 - 1:200 - Santa Cruz<sup>®</sup> |
| Oxphos | Cocktail ab123545 - 1:1000 - Abcam<sup>®</sup> |

The membranes were then incubated with secondary antibody (1:10000, Santa Cruz®), under agitation, followed by development with chemiluminescent reagents (Thermo Scientific®) and photodocumentation using G:BOX (Syngene®). The intensities of the bands were determined by densitometry, using ImageJ® software (NIH, USA).

### Histological Analysis of Liver Tissue

After fixation in 10% formalin solution for twenty for four hours, small fragments of liver tissue were embedded in paraffin using usual procedures. Sections of 4μm thick with 20μm of distance between the sections were stained with hematoxylin-eosin solution for histological analysis. The images were captured by a Leica DM 2000® photomicroscope with Las software (version 4.1®). Ten images were captured per animal using the 400X magnification analyzed by Image-Pro Plus® (version 4.5.0.29). Stereological Analysis: In each of the ten images, a grid was applied with three hundred intersections totaling three thousand intersections per animal. By image, the intersections were classified into hepatocyte cytoplasm, mononuclei, binuclei, lipid deposit and connective tissue elements (vessels, connective cells and matrix). The frequencies (%) of each element are presented as mean ± SEM. Cell Quantification and Morphometry: In each of the ten images, all hepatocytes present in the area were counted. Only fully delimited cells were considered. Morphometry involved defining the diameter of the hepatocyte and its nucleus in a total of thirty cells per animal. Values for all cells are presented as mean ± SEM.

### Histological analysis of the pancreas

After fixation of the pancreas, small fragments were embedded in paraffin using standard procedures, with dehydration in increasing concentrations of alcohol. Sections (5μm thick) were stained with hematoxylin-eosin solution, Perls (for analysis of iron deposits), and Gomori’s trichrome (for analysis of collagen fibers). Two sections were used for each SHAM, PH, and OHR animal, totaling 12 sections per animal group. Images were acquired using a Leica DM 2000® photomicroscope operated with LAS v.4.1® software.

Qualitative histological evaluations of the Perls and Gomori staining results were performed by two observers, considering the presence and absence of staining. The evaluations included the entire area of the section, employing images acquired at 400x magnification, which were analyzed using Image-Pro Plus® (v.4.5.0.29) software. For each animal, the diameters were obtained for all the islets that were fully delimited in the image.

### Detection of IDE mRNA by RT-PCR

Total RNA was isolated from approximately 100 mg of rat liver using TRIzol® reagent (Invitrogen®, Carlsbad, CA, USA), including the digestion of contaminating DNA with amplification grade DNAse I (Invitrogen®), following the manufacturer’s instructions. RNA purity and concentration were determined spectrophotometrically (Thermo Fisher Scientific Evolution™ 249 300 UV-Vis) in OD_260_ for concentration and purity in OD_260_/OD_280_ = above 1,6. Synthesis of cDNA employed 2µg of RNA in the presence of dithiothreitol, dNTP, random primers, RNAseOUT, and SuperScript™ II Reverse Transcriptase (Invitrogen®), in a final volume of 20µL. The mRNA levels of the insulin-degrading enzyme (IDE) genes were investigated by semi-quantitative RT-PCR. The primer sequences used in the PCR reactions were chosen based on the sequences available in GenBank. IDE gene amplification was performed sense 5′-AGGAAATGTTGGCTGTGGACGCA-3′ and antisense 5′-CCTGGCAAGAACGTGGACGGATA-3′ primers to amplify a predicted amplicon of 62bp (Tm 57 °C). The amplified products were separated on 2.0% agarose gel stained with ethidium bromide and the gel was then photographed using LPix-Touch® (Loccus Biotecnologia®). The signal intensities of the bands were obtained using G:Box (Syngene®), with densitometric measurement using Scion Image® software (Scion Corp., Frederick, MD, USA). Each value was obtained as the mean of three densitometry readings. The results were expressed as average ratios of the relative expression of transcripts normalized with β-actin as the control housekeeping gene. β-actin gene amplification was performed using gene forward AGAGGGAAATCGTGCGTGACA and reverse CGATAGTGATGACCTGACCGTA primers to amplify a predicted amplicon of 138bp (Tm 57 °C).

### Different networks of animal markers stratified into firm groups

Data from marker networks in obese and hypertensive animals, diagnosed or not with restricted dietary, together with SHAM using special Spearman correlations at p<0.05. From the analysis of the correlation of all groups, the correlations were performed using the Cytoscape® 3.0.4 software (Cytoscape Consortium San Diego, CA, USA). The data were represented by connecting edges to highlight positive (strong r≥0.68; thick continuous line), moderate (0.36≥ r ≤0.68; thinner continuous line) or weak (r <0.35; thin continuous line) and negative (r ≤-0.68; thick connected line), moderate (−0.68≥ r ≤-0.36; thinner initiated line) or weak (r >-0.36; initiated line thin) as previously proposed [1]. The absence of the line indicates the absence of the relationship.

### Statistical analysis

For comparison of multiple groups, we performed one-way analysis of variance (ANOVA) followed by Tuckey post-test. The differences between any two groups were evaluated using two-tailed Student’s t-test. All calculations were performed in the GraphPad Prism 9.0 software (GraphPad®, San Diego, CA, USA). Differences with P<0.05 were considered statistically significant. The results were expressed as mean ± standard error of the mean (X±SEM).

## Results

### Caloric restriction promotes reestablishment of blood pressure, body weight and recovers glycemic homeostasis in obese and hypertensive animals

After being allocated and paired, the animals were monitored throughout the experiment, the groups and the different dietary interventions are represented in a graphic summary (**Figure 1A**). Rats were submitted to renovascular arterial hypertension using the Goldblatt 2K1C technique and they were put on a diet that was high in fat (HFD) or remained on a normal chow diet. Following 10 wk on an HFD, a subset of the HFD group underwent CR for 4 wk (OHR). After 10 wk, the HFD feeding induced a significant increase in the body weight and Lee index of mice compared with the SHAM and OHR groups **(Fig. 1B and F)**. OHR promoted a significant reduction in the body weight and blood pressure compared with the OH group **(Figure 1B and C)**, in agreement with the literature [10, 15]. In **Figure 1D**, despite being offered *ad libitum*, ingestion of the diet rich in lipids was lower, suggesting that it provided satiety after consumption of a smaller quantity and/or that the animals of the OH group did not find it pleasant. Despite the decreased ingestion, the animals that received the lipid-rich feed showed increases of body weight and the Lee index **(Figure 1B and F)**, due to lipid/calorie overload and nutritional imbalance of the food. In the OHR group, despite the excessive lipid load, the animals consumed less food due to the CR, so body weight was similar to that of the SHAM group, as reported previously [10]. There was no difference in serum concentrations of proteins **(data not show)**. As expected [11], the CR decreases serum glucose and improve de lipid profile **(Figure 1E and Figure 2A, C, D, E, I)**. In **Figure 1G**, the GTT and area under the curve values revealed intolerance to glucose in the OH group and reestablishment of control in the OHR group, when compared to the SHAM group (OH: 26409±2039; OHR: 22950±291.2; SHAM: 19766±1467). In **Figure 1H**, the ITT indicated greater insulin sensitivity in the OHR group, compared to the SHAM and OH groups, which was reflected in the Kitt values shown in **Figure 1H** (OHR: 1.58±0.14 %/min; SHAM: 1.24±0.09 %/min; OH: 0.78±0.10 %/min). HDL (**Figure 2B**) and hepatic cholesterol (**Figure 2H**) were both significantly elevated in CR group. **Figure 2J** shows that there was increased IDE gene expression in the livers of group OH, compared to the SHAM group (1.2±0.09 and 0.81±0.04, respectively), and similarity between the OHR and SHAM groups (0.92±0.06 and 0.81±0.04, respectively). The IDE can preserve the means for peripheral insulin sensitivity, according to our results obtained in GTT and ITT. **Figure 2F** shows that the hepatic glycogen level (in g per 100g of tissue) was lower in the OHR group, compared to the SHAM group, (0.32±0.045 and 0.52±0.04, respectively), while levels were similar for the OH and SHAM groups. Reduced glycogen stores in OHR animals strengthen adaptive physiological mechanisms of weight loss. CR animals initially have reduced or depleted glycogen stores, together with decreased insulin secretion, both of which are required in order to meet the energy needs of the brain [23]. There was no difference in muscle glycogen (**Figure 2G)** for the all groups.

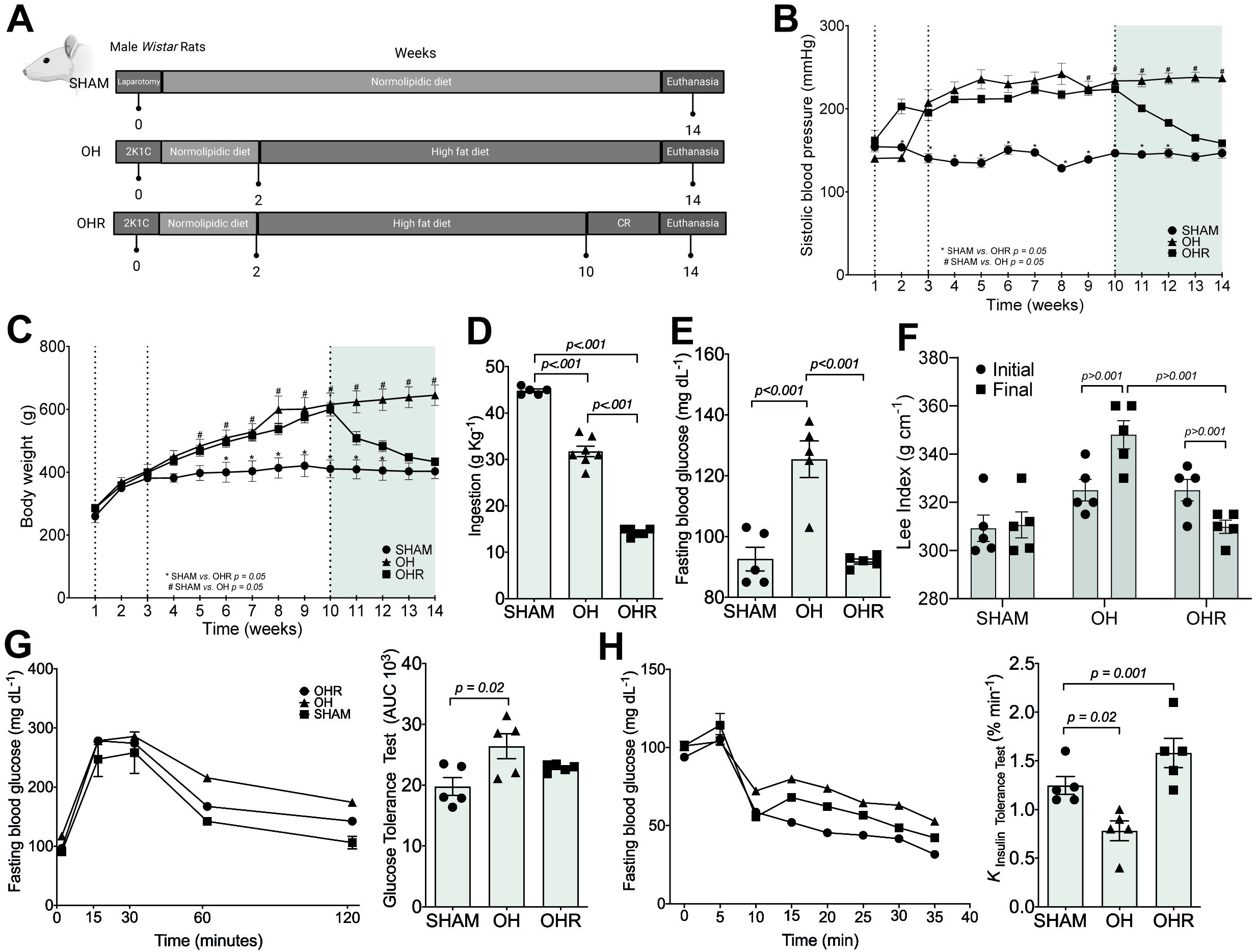

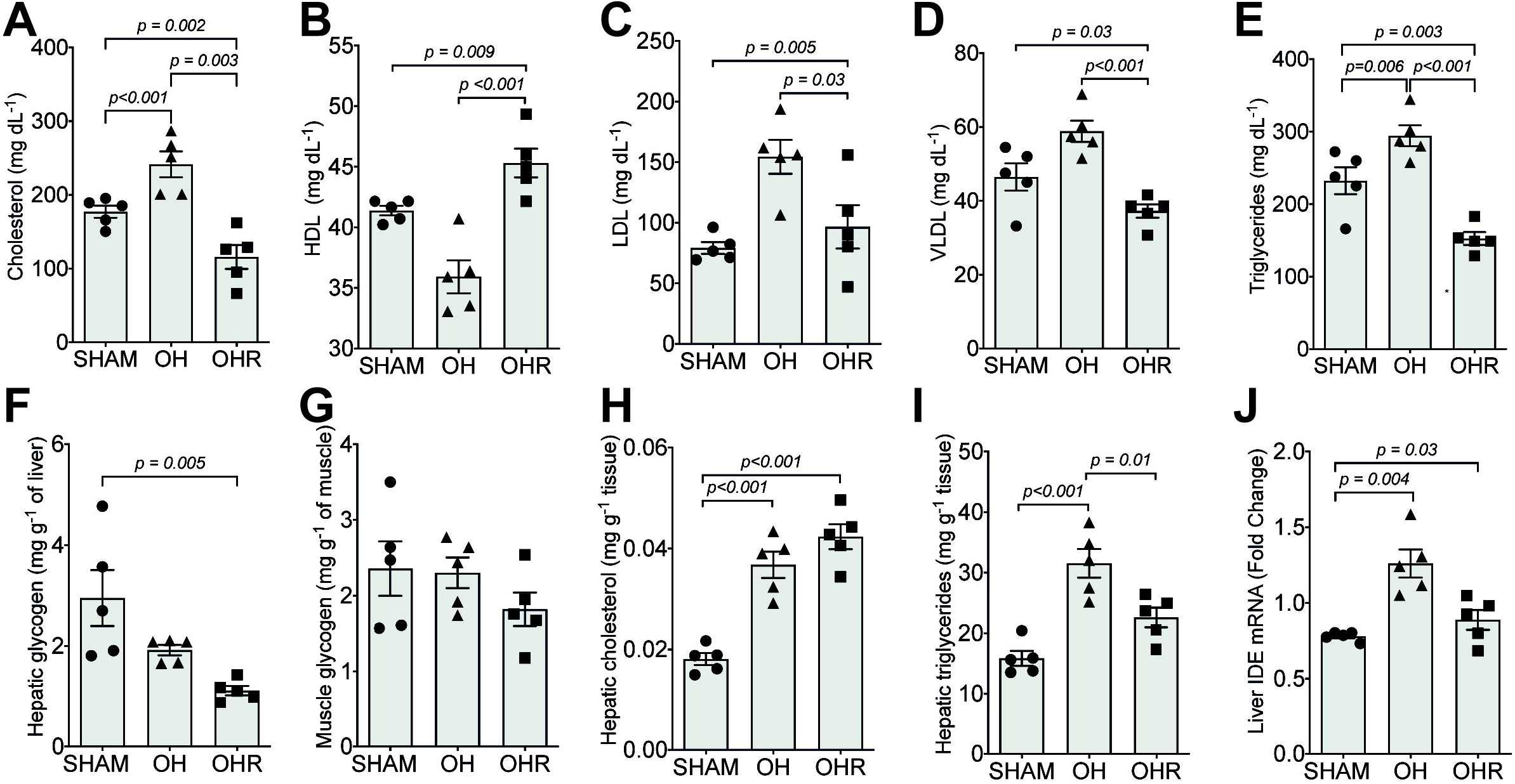

### Caloric restriction preserves pancreatic structures, reverses islet fibrosis, but causes an increase in lipid drops in hepatocytes

In **Figure 3A**, representative images from liver and pancreas it can be seen that the slides stained by the H&E, Perls and Trichrome Gomori technique. No differences were observed for aspartate aminotransferase, alanine aminotransferase levels **(Figure 3 B and C)** and for the diameters of the islets from H&E technique **(Figure 3F)**. Perls technique **(Figure 3G)** showed an absence of iron deposits (hemosiderin) for the three groups (SHAM, OH, and OHR). The slides stained using Gomori’s trichrome **(Figure 3H)** showed that there was higher incidence of collagen in the islets of the OH group, compared to the OHR and SHAM groups. The fibrotic pancreatic islet has been shown in human type 2 diabetes, as well as in rodent models [6,8]. Our results showed that CR contribute to islet functional preservation characterized by absent of collagen associated an increased the insulin levels **(Figure 3 E)**. According to the histological data, Table 3, the diameter of the hepatocytes, area occupied by conjunctive element and binuclear and mononuclear cells were similar in all animal groups and suggests cellular integrity. The increase in the area of the hepatocyte nucleus in the OHR group (23.81%) compared to SHAM (4.76%) and OH (4.79%) may indicate elevated hepatic proliferative capacity, which is an organ presenting high degree of cell regeneration. Thus, that is suggest that nucleus occupy space in the hepatocyte cytoplasm that were reduced in the OH group (26.54%) and OHR (26.82%) versus SHAM (46.19%). Hepatocytes showed no signs of inflammation and fibrosis in any of the groups suggesting hepatic integrity.

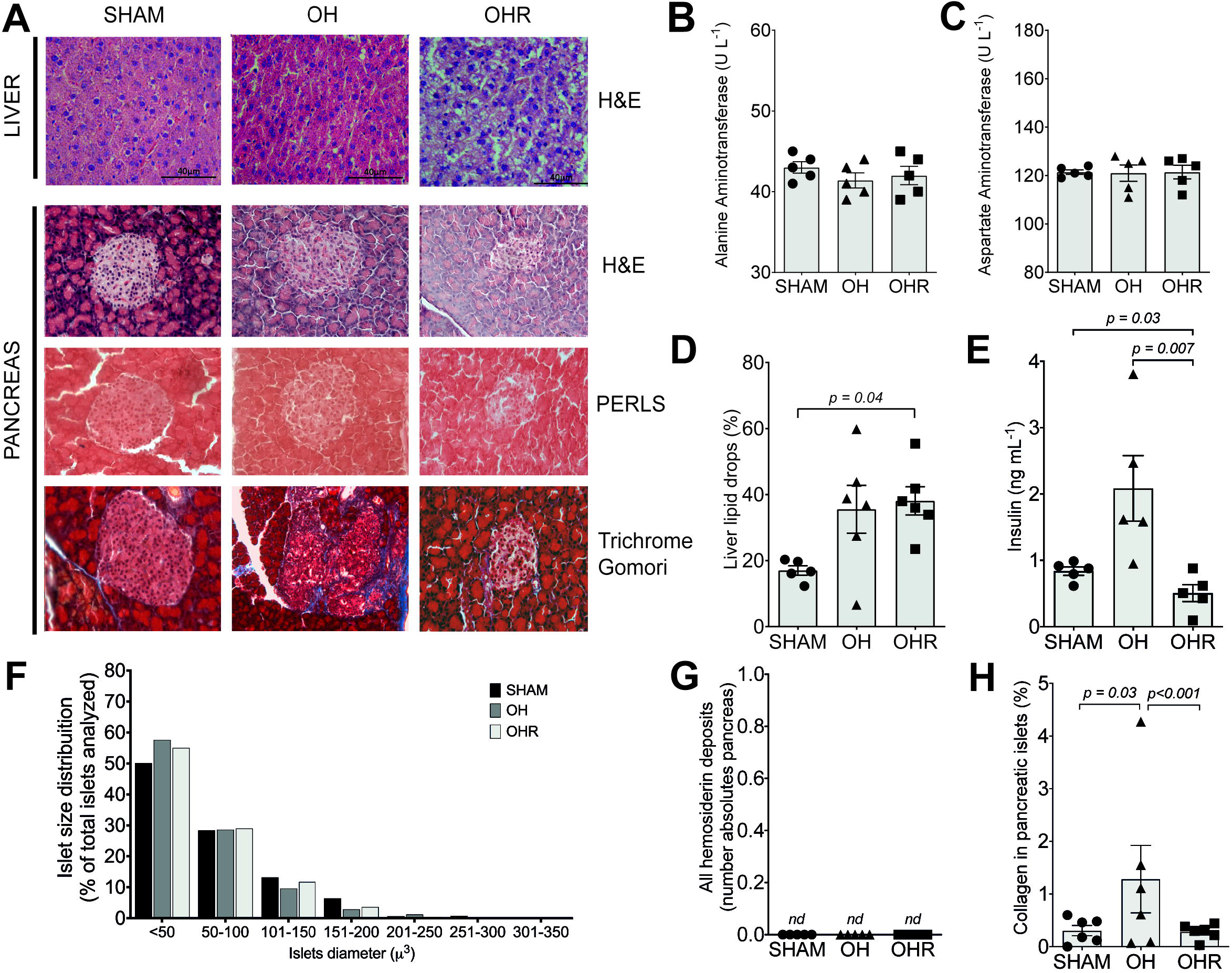

**Table 3.** Stereological and morphometric profiles of the hepatic tissues of the Sham, HO, and HOR group animals.

| Parameters | Sham | HO | HOR |
| --- | --- | --- | --- |
| Mononucleate (%) | 18.55 | 17.58 | 17.78 |
| Binucleate (%) | 4.45 | 5.27 | 5.84 |
| Lipid (%) | 16.99 | 35.54* | 38.09** |
| Cytoplasm (%) | 46.19 | 26.54* | 26.82** |
| Conjunctive element (%) | 13.49 | 13.65 | 11.38 |
| Number of cells (n) | 183 | 192 | 226 |
| Cell nucleus diameter | 4.76 | 4.79 | 23.81**<sup>+</sup> |
| Hepatic cell diameter (μm) | 13.53 | 13.90 | 13.61 |

ANOVA with the Tukey post-test: \*p<0.05 Sham *versus* HO; \*\*p<0.05 Sham *versus* HOR; <sup>+</sup>p<0.05 HO *versus* HOR; mean ± SEM (n=6).

### Oxidative stress is reversed during caloric restriction mediated by SIRT1, SIRT3 and Nampt, establishing a strong correlation with lipid metabolism

As shown in **Figure 4A**, there was higher expression of insulin receptor (IR) in the OHR group (14.92±0.53), compared to the SHAM group (4.64±1.98) and OH group (3.81±1.30). The expression of SIRT1 (**Figure 4D**) was higher for the OHR group (7.02±0.88) compared to the SHAM group (4.48±0.91) and OH group (3.24±0.87). Nampt was higher for the OHR group (12.62±0.87) compared to the SHAM (3.34±0.32) and OH group (2.61±0.34) (**Figure 4G**). IR and SIRT1 acts in the regulation of glucose metabolism and reflect in improve of GTT and ITT of OHR group. Given that Nampt is directly regulates the activity of SIRT1 our data seems to establish metabolic hepatic regulation in OHR. The values obtained for the other proteins **(Figure B, C, E, F, H)** were similar for all the groups. **Figure 4** I shows that the expression of complex II was low in the OH group (5.355±0.79) and was significantly higher in the OHR group (11.08±1.53). Given the importance of mitochondrial complex II [2,13] in the maintenance of cellular integrity, probably that our results indicate mitochondrial modifications due to diet-related by preserve the integrity of OXPHOS in the livers of obese/ hypertensive/restricted animals.

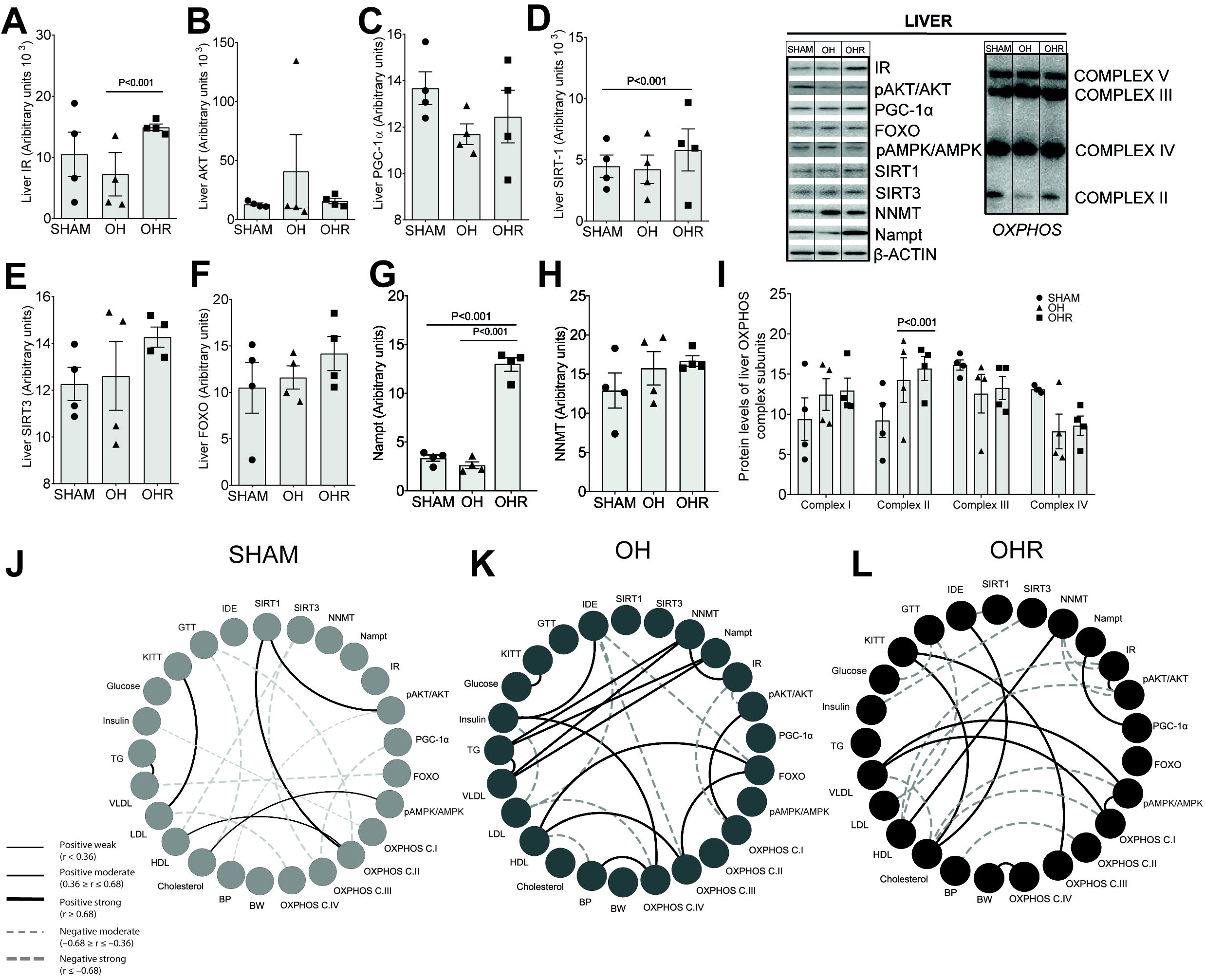

In **figure 4 J, K and L** correlation data. The degree of significance is represented by the thickness of the line. Correlations were created using the Spearman test; p values were used to classify as weak (r≤0.35, p<0.05), moderate (r=0.36–0.67, p<0.01) or strong (r ≥0.68, p<0.001). The absence of the line indicates the absence of the relationship. To assess the relationship between the markers, SIRT1, SIRT3, NNMT, Nampt, IR, pAKT/AKT, PCG-1α, FOXO, pAMPK/AMPK, OXPHOS, Body Weight (BW), Blood Pressure (BP), Cholesterol, HDL, LDL, VLDL, Triglycerides, Insulin, Glucose, kITT, GTT and IDE, a series of correlation analyzes was performed **(Figure 4 J, K, L)**. When comparing the interactions analyzed between the SHAM group profile, OH and OHR. In SHAM (J), we observed that interactions occur mainly in SIRT1 with pAKT/AKT (moderate), SIRT1 with OXPHOS C.II (moderate), OXPHOS C.II and HDL, cholesterol between pAMPK/AMPK. LDL and kITT have a moderate interaction, as do VLDL and triglycerides. In OH (K), SIRT1 and SIRT3 had no interaction with the other studied markers. NNMT was positively related in a moderate way to triglycerides, VLDL and IR; Nampt, in a positive and moderate way, correlated with triglycerides and VLDL; pAKT/AKT there was a positive and moderate correlation with OXPHOS C.I, as well as FOXO with HDL and OXPHOS C.III; Insulin and Blood Pressure with C.IV, IDE with insulin and glucose, as expected, correlated positively and moderately with kITT. We detected a positive and moderate correlation between VLDL and Triglycerides, as expected. In the animals of the OHR group (L), there was a moderate interaction between SIRT1 and IDE as well as between SIRT3 and Insulin; NNMT interacted moderately and positively with HDL, but moderately with IR and pAKT/AKT; Nampt interacted moderately and positively with PGC-1α; pAMPK/AMPK, moderately and positively interacted with VLDL and OXPHOS C.I, but negatively with cholesterol. C.II interacted negatively with blood pressure, however C.III had an interaction of the same intensity but positively with kITT, as well as C.IV and its interaction with the animals’ body weight. Cholesterol had a moderate and positive interaction with kITT and IDE. Finally, glucose in a negative way with GTT.

## Discussion

The glucose and lipid homeostasis requires continuous control to blood sugar level and body weight to minimize the risks of metabolic dysfunctions. Thus, pancreatic islets and liver are critical tissues which is involved in this control. Pancreatic islets released insulin; it stimulates the entrance of glucose into skeletal muscles, liver and adipose tissue via special transporters, thus controlling glucose homeostasis. Already the removal of insulin from the physiological system is dependent on the IDE present in various tissues, predominantly the liver [33]. OH rat’s group developed overweight, hypertension and pre diabetes, as shown by glucose intolerance, reduction of glycogen stores, peripheral insulin resistance, mild hyperglycemia and hyperinsulinemia, increase in hepatic IDE and increase of pancreatic of collagen areas. Islet inflammation in the type 2 diabetes likely is the cause of the fibrosis and collagen deposit [6]. HFD feeding previously reported to increase of macrophages in islets of nonhuman primates [24]. In fact, the injured islets of the HFD rats showed similar fibrosis and suggests possible link between pre diabetic features. The deficit of glucose homeostasis of the OH animals, together with the increase of hepatic IDE, was suggestive of imbalance of insulin clearance and corroborates with our correlation study. Previously was found in animals submitted to a cafeteria diet [3] incresead IDE activity in the liver, corroborating the present results [4]. The hyperlipidic diet with 40% CR for four weeks reestablished control of glucose metabolism by preventing pancreatic fibrosis, ensuring insulin sensitivity and reduction of hepatic IDE, while at the same time reducing arterial blood pressure [16]. The pressure recovery observed in the OHR group is in agreement with the literature showing that the administration of NAD+ in mice or primary cardiomyocytes exposed to hypertrophic agonists led to improvement of the hypertrophic phenotype [26,27]. CR has been shown to reverse endothelial alterations in a diet-induced obesity model reduce vascular oxidative stress in OLETF (Otsuka Long Evans Tokushima Fatty) rats, and reduce arterial blood pressure in spontaneously hypertensive rats [15]. As previously reported in a study when were reduced mRNA expression of IDE could provide an explanation for the reduced of serum insulin in group OHR [32]. This mechanism acted to increase the peripheral insulin sensitivity through increased expression of IR observed in the liver, suggesting that it was associated with decreased insulin clearance. Our findings suggest that hepatic mechanism for these physiological changes, OHR group, was increased expression of the hepatic proteins IR and Nampt, with subsequent increase of SIRT1 and complex II of the electron transport chain, as well as mitochondrial adaptive of OXPHOS. OHR group the CR led to activation of SIRT1 by Nampt, as reported previously [5]. Also was reported that after partial withdrawal of food, the SIRT1 protein binds to and represses genes responsible for fat storage, with its overexpression attenuating adipogenesis and its upregulation leading to lipolysis [25]. Due to the food restriction, it is likely that fat provided an energy source for the maintenance of cellular metabolism. SIRT1 provides protection against diseases related to insulin resistance, such as the metabolic syndrome and diabetes mellitus [19, 29]. In mice, the general knockout of Nampt resulted in lethality in homozygotes, while heterozygotes presented glucose intolerance [28]. Other studies found that Nampt mRNA expression levels decreased in the livers of obese mice fed a diet rich in fat [35]. On the other hand, fasting was found to increase hepatic Nampt levels, NAD+ biosynthesis, and hepatic metabolic function, leading to reduced accumulation of triglycerides [34,35]. In contrast, transgenic mice with enzymatically inactivated Nampt presented increased accumulation of triglycerides, insulin resistance, and a phenotype of non-alcoholic fatty liver disease (NAFLD) [31]. An important finding was that the NNMT protein showed similarity among the groups (SHAM, OH, and OHR), since this protein can inhibit SIRT1 [18]. However, a previous study suggested that increased NNMT expression could stabilize the SIRT1 protein, resulting in metabolic benefits [14]. Meanwhile, knockdown of NNMT in white adipose tissue and the liver can protect against obesity induced by diet [18]. Modulation of complex II in the OH and OHR animals was suggestive of mitochondrial adaptive, as a function of the imposed diets [23]. The Akt, AMPK, PGC1α, and FoxO3 proteins showed no alterations in the groups studied, despite the benefits reported in the literature [22]. In an addition, in our correlation study the animals in the OHR group point to a strong interaction of IR with AKT, Nampt and PGC1 and kITT and lipid profile and OXPHOS C.III. Also there was correlation OXPHOS C.I with VLDL and pAMPK/AMPK and body weight with OXPHOs C.IV corroborating with our suggestion that caloric restriction recovers the pre-diabetic metabolic profile of the submitted hypertensive animal the fat-rich diet. It is possible that if the CR was modified, with longer duration and/or greater severity, the expression of these proteins might be increased, hence affecting the metabolism. Studies concerning CR vary considerably in terms of duration, animal age, and diet adopted, but a commonly used protocol is a 40% restricted diet (moderate CR), compared to the control, for four weeks [10].

## Conclusion

It could be concluded that short-term CR provided benefits in terms of glucose and lipids homeostasis, with protection of the pancreatic islets and hepatic cells against fibrosis, as well as improved arterial blood pressure in obese and hypertensive animals. Although our rodent models of obese/hypertension do not always perfectly reproduce the human pathology of obesity and hypertension, it has helped to better our knowledge of the cell, metabolic and molecular pathways of lipid and glucose homeostasis. Critical proteins activated by CR like the one Nampt, SIRT1 and OXPHOS have been involved in many metabolic defects associated with obesity and hypertension. Therefore, a better understanding of the regulation of the proteins that control the effects of CR may assist in the development of therapeutic approaches for this metabolic disease.

## Author’s contribution

MECA, CAO, BFT conceived and carried out the experimental design of the project. CAO performed renal artery stenosis to induce hypertension. VEP, MCB, JVH and NTH carried out dietary interventions and observations throughout the period. VEP, MCB, JVH, NTH and BFT performed the processing and histological analysis. VEP and MSSNF performed the biochemical tests. CAO, VEP, MSSNF and MECA performed the molecular assays. PVSN and VEP performed the study of correlations and built the figures. MECA, VEP and MSSNF edited the final manuscript. All authors read and approved the final version of the manuscript.

## Acknowledgments

The authors are grateful Edvaldo Costa, BSc; Sr Mateus Eduardo Bortolanza da Silva; Sra Ana Cristina Pires Menegheti; and Renata Barbieri for their excellent technical assistance.

## Additional information

Available in the Supplementary Appendix

