## Supplementary Appendix for "Caloric restriction overcomes pre-diabetic and hypertension induced by high fat diet and renal artery stenosis"

### Networks SHAM Additional data

| SHAM | SIRT1 | SIRT3 | NNMT | Nampt | IR | pAKT/AKT | PGC-1a | FOXO | AMPK/AM | HOS Comp | HOS Comp | HOS Comp | HOS Compl | PC | PA | Colesterol | HDL | LDL | VLDL | Triglicérides | Insulina | Glicemia | KITT | GTT | IDE |
|---|---|---|---|---|---|---|---|---|---|---|---|---|---|---|---|---|---|---|---|---|---|---|---|---|---|
| SIRT1 | | | | | | | | | | | | | | | | | | | | | | | | | |
| SIRT3 | 0.75 | | | | | | | | | | | | | | | | | | | | | | | | |
| NNMT | 0.333333 | 0.916667 | | | | | | | | | | | | | | | | | | | | | | | |
| Nampt | 0.416667 | 0.75 | 0.333333 | | | | | | | | | | | | | | | | | | | | | | |
| IR | 0.916667 | 1 | 0.75 | 0.916667 | | | | | | | | | | | | | | | | | | | | | |
| pAKT/AKT | 0.083333 | 0.75 | 0.333333 | 0.416667 | 0.916667 | | | | | | | | | | | | | | | | | | | | |
| PGC-1a | 1 | 0.333333 | 0.75 | 0.333333 | 0.75 | 1 | | | | | | | | | | | | | | | | | | | |
| FOXO | 0.75 | 0.75 | 1 | 0.75 | 0.333333 | 0.75 | 0.916667 | | | | | | | | | | | | | | | | | | |
| pAMPK/AMP | 0.333333 | 0.333333 | 0.75 | 1 | 0.75 | 0.333333 | 0.416667 | 0.916667 | | | | | | | | | | | | | | | | | |
| OXPHOS Complex I | 0.333333 | 0.916667 | 0.416667 | 0.333333 | 0.75 | 0.333333 | 0.75 | 0.333333 | 0.75 | | | | | | | | | | | | | | | | |
| OXPHOS Complex II | 0.083333 | 0.75 | 0.333333 | 0.416667 | 0.916667 | 0.083333 | 1 | 0.75 | 0.333333 | 0.333333 | | | | | | | | | | | | | | | |
| OXPHOS Complex III | 0.75 | 0.083333 | 0.916667 | 0.75 | 1 | 0.75 | 0.333333 | 0.75 | 0.333333 | 0.916667 | 0.75 | | | | | | | | | | | | | | |
| OXPHOS Complex IV | 0.75 | 0.416667 | 0.916667 | 0.75 | 0.333333 | 0.75 | 0.333333 | 0.75 | 0.333333 | 0.916667 | 0.75 | 0.416667 | | | | | | | | | | | | | |
| PC | 0.916667 | 1 | 0.75 | 0.916667 | 0.083333 | 0.916667 | 0.75 | 0.333333 | 0.75 | 0.75 | 0.916667 | 1 | 0.333333 | | | | | | | | | | | | |
| PA | 0.75 | 0.75 | 1 | 0.75 | 0.333333 | 0.75 | 0.916667 | 0.083333 | 0.916667 | 0.333333 | 0.75 | 0.75 | 0.75 | 0.233333 | | | | | | | | | | | |
| Colesterol | 0.333333 | 0.916667 | 0.416667 | 0.333333 | 0.75 | 0.333333 | 0.75 | 0.333333 | 0.75 | 0.083333 | 0.333333 | 0.916667 | 0.916667 | 0.95 | 0.45 | | | | | | | | | | |
| HDL | 0.75 | 0.083333 | 0.916667 | 0.75 | 1 | 0.75 | 0.333333 | 0.75 | 0.333333 | 0.916667 | 0.75 | 0.083333 | 0.416667 | 0.833333 | 0.266667 | 0.733333 | | | | | | | | | |
| LDL | 1 | 0.333333 | 0.75 | 0.333333 | 0.75 | 1 | 0.083333 | 0.916667 | 0.416667 | 0.75 | 1 | 0.333333 | 0.333333 | 0.45 | 0.95 | 0.683333 | 0.166667 | | | | | | | | |
| VLDL | 0.916667 | 0.333333 | 0.75 | 0.916667 | 0.416667 | 0.916667 | 0.75 | 0.333333 | 0.75 | 0.75 | 0.916667 | 0.333333 | 1 | 0.95 | 0.95 | 0.783333 | 0.9 | 0.683333 | | | | | | | |
| Triglicérides | 0.916667 | 0.333333 | 0.75 | 0.916667 | 0.416667 | 0.916667 | 0.75 | 0.333333 | 0.75 | 0.75 | 0.916667 | 0.333333 | 1 | 0.95 | 0.95 | 0.783333 | 0.9 | 0.683333 | 0.016667 | | | | | | |
| Insulina | 0.333333 | 0.333333 | 0.75 | 1 | 0.75 | 0.333333 | 0.416667 | 0.916667 | 0.083333 | 0.75 | 0.333333 | 0.333333 | 0.333333 | 0.683333 | 1 | 0.45 | 0.266667 | 0.35 | 0.95 | 0.95 | | | | | |
| Glicemia | 1 | 0.833333 | 0.5 | 0.666667 | 0.166667 | 1 | 0.666667 | 0.333333 | 1 | 0.833333 | 1 | 0.833333 | 0.5 | 0.1 | 0.4 | 0.833333 | 0.666667 | 0.733333 | 0.4 | 0.4 | 0.733333 | | | | |
| KITT | 0.833333 | 0.166667 | 1 | 0.5 | 0.833333 | 0.833333 | 0.166667 | 0.666667 | 0.333333 | 0.666667 | 0.833333 | 0.166667 | 0.333333 | 0.566667 | 0.833333 | 0.733333 | 0.15 | 0.033333 | 0.5 | 0.5 | 0.266667 | 1 | | | |
| GTT | 0.333333 | 0.916667 | 0.416667 | 0.333333 | 0.75 | 0.333333 | 0.75 | 0.333333 | 0.75 | 0.083333 | 0.333333 | 0.916667 | 0.916667 | 0.783333 | 0.683333 | 0.083333 | 1 | 0.95 | 0.45 | 0.45 | 0.233333 | 0.566667 | 1 | | |
| IDE | 0.75 | 0.75 | 0.333333 | 0.75 | 0.333333 | 0.75 | 0.916667 | 0.416667 | 0.916667 | 1 | 0.75 | 0.75 | 0.75 | 0.1 | 0.166667 | 0.833333 | 0.433333 | 0.733333 | 1 | 1 | 0.733333 | 0.15 | 1 | 0.833333 | |

### Networks OH Additional data

| OH | SIRT1 | SIRT3 | NNMT | Nampt | IR | pAKT/AKT | PGC-1a | FOXO | AMPK/AM | HOS Comp | HOS Comp | HOS Comp | IOS Compl | PC | PA | Colesterol | HDL | LDL | VLDL | Triglicérides | Insulina | Glicemia | KITT | GTT | IDE |
|---|---|---|---|---|---|---|---|---|---|---|---|---|---|---|---|---|---|---|---|---|---|---|---|---|---|
| SIRT1 | | | | | | | | | | | | | | | | | | | | | | | | | |
| SIRT3 | 0.75 | | | | | | | | | | | | | | | | | | | | | | | | |
| NNMT | 0.416667 | 0.75 | | | | | | | | | | | | | | | | | | | | | | | |
| Nampt | 0.416667 | 0.75 | 0.083333 | | | | | | | | | | | | | | | | | | | | | | |
| IR | 1 | 0.333333 | 0.333333 | 0.333333 | | | | | | | | | | | | | | | | | | | | | |
| pAKT/AKT | 1 | 0.333333 | 0.333333 | 0.333333 | 0.083333 | | | | | | | | | | | | | | | | | | | | |
| PGC-1a | 0.75 | 0.75 | 0.75 | 0.75 | 0.916667 | 0.916667 | | | | | | | | | | | | | | | | | | | |
| FOXO | 0.916667 | 1 | 0.916667 | 0.916667 | 0.75 | 0.75 | 0.333333 | | | | | | | | | | | | | | | | | | |
| pAMPK/AMP | 0.75 | 0.416667 | 0.75 | 0.75 | 0.333333 | 0.333333 | 0.75 | 0.333333 | | | | | | | | | | | | | | | | | |
| OXPHOS Complex I | 1 | 0.333333 | 0.333333 | 0.333333 | 0.083333 | 0.083333 | 0.916667 | 0.75 | 0.333333 | | | | | | | | | | | | | | | | |
| OXPHOS Complex II | 0.333333 | 0.333333 | 1 | 1 | 0.416667 | 0.416667 | 0.916667 | 0.75 | 0.333333 | 0.416667 | | | | | | | | | | | | | | | |
| OXPHOS Complex III | 0.916667 | 1 | 0.916667 | 0.916667 | 0.75 | 0.75 | 0.333333 | 0.083333 | 0.333333 | 0.75 | 0.75 | | | | | | | | | | | | | | |
| OXPHOS Complex IV | 0.75 | 0.75 | 0.75 | 0.75 | 0.916667 | 0.916667 | 0.416667 | 0.333333 | 0.75 | 0.916667 | 0.916667 | 0.333333 | | | | | | | | | | | | | |
| PC | 0.916667 | 0.333333 | 0.916667 | 0.916667 | 0.75 | 0.75 | 0.333333 | 0.416667 | 1 | 0.75 | 0.75 | 0.416667 | 0.333333 | | | | | | | | | | | | |
| PA | 0.75 | 0.75 | 0.75 | 0.75 | 0.916667 | 0.916667 | 0.416667 | 0.333333 | 0.75 | 0.916667 | 0.916667 | 0.333333 | 0.083333 | 0.233333 | | | | | | | | | | | |
| Colesterol | 0.166667 | 1 | 0.333333 | 0.333333 | 0.833333 | 0.833333 | 0.833333 | 0.666667 | 0.666667 | 0.833333 | 0.5 | 0.666667 | 0.5 | 0.9 | 0.166667 | | | | | | | | | | |
| HDL | 0.916667 | 1 | 0.916667 | 0.916667 | 0.75 | 0.75 | 0.333333 | 0.083333 | 0.333333 | 0.75 | 0.75 | 0.083333 | 0.333333 | 0.35 | 0.083333 | 0.4 | | | | | | | | | |
| LDL | 0.75 | 0.75 | 0.75 | 0.75 | 0.916667 | 0.916667 | 0.416667 | 0.333333 | 0.75 | 0.916667 | 0.916667 | 0.333333 | 0.083333 | 0.233333 | 0.233333 | 0.4 | 0.35 | | | | | | | | |
| VLDL | 0.416667 | 0.75 | 0.083333 | 0.083333 | 0.333333 | 0.333333 | 0.75 | 0.916667 | 0.75 | 0.333333 | 1 | 0.916667 | 0.75 | 0.783333 | 0.683333 | 0.833333 | 0.35 | 0.783333 | | | | | | | |
| Triglicérides | 0.416667 | 0.75 | 0.083333 | 0.083333 | 0.333333 | 0.333333 | 0.75 | 0.916667 | 0.75 | 0.333333 | 1 | 0.916667 | 0.75 | 0.783333 | 0.683333 | 0.833333 | 0.35 | 0.783333 | 0.016667 | | | | | | |
| Insulina | 0.75 | 0.75 | 0.75 | 0.75 | 0.916667 | 0.916667 | 0.416667 | 0.333333 | 0.75 | 0.916667 | 0.916667 | 0.333333 | 0.083333 | 0.233333 | 0.233333 | 0.4 | 0.35 | 0.016667 | 0.783333 | 0.783333 | | | | | |
| Glicemia | 0.333333 | 0.916667 | 0.333333 | 0.333333 | 0.75 | 0.75 | 1 | 0.75 | 0.916667 | 0.75 | 0.75 | 0.75 | 0.333333 | 0.783333 | 0.683333 | 0.3 | 1 | 0.233333 | 0.133333 | 0.133333 | 0.233333 | | | | |
| KITT | 0.166667 | 0.5 | 1 | 1 | 1 | 1 | 1 | 1 | 0.5 | 1 | 0.166667 | 1 | 1 | 1 | 0.266667 | 0.1 | 0.266667 | 0.833333 | 0.633333 | 0.633333 | 0.833333 | 1 | | | |
| GTT | 0.333333 | 0.916667 | 0.333333 | 0.333333 | 0.75 | 0.75 | 1 | 0.75 | 0.916667 | 0.75 | 0.75 | 0.75 | 0.333333 | 0.783333 | 0.683333 | 0.3 | 1 | 0.233333 | 0.133333 | 0.133333 | 0.233333 | 0.016667 | 1 | | |
| IDE | 0.916667 | 1 | 0.916667 | 0.916667 | 0.75 | 0.75 | 0.333333 | 0.083333 | 0.333333 | 0.75 | 0.75 | 0.083333 | 0.333333 | 0.35 | 0.35 | 0.5 | 0.233333 | 0.083333 | 0.95 | 0.95 | 0.083333 | 0.516667 | 0.833333 | 0.516667 | |

### Networks OHR Additional data

| OHR | SIRT1 | SIRT3 | NNMT | Nampt | IR | pAKT/AKT | PGC-1a | FOXO | AMPK/AM | HOS Comp | HOS Comp | HOS Comp | HOS Compl | PC | PA | Colesterol | HDL | LDL | VLDL | Triglicérides | Insulina | Glicemia | KITT | GTT |
|---|---|---|---|---|---|---|---|---|---|---|---|---|---|---|---|---|---|---|---|---|---|---|---|---|
| SIRT1 | | | | | | | | | | | | | | | | | | | | | | | | |
| SIRT3 | 0.416667 | | | | | | | | | | | | | | | | | | | | | | | |
| NNMT | 1 | 0.333333 | | | | | | | | | | | | | | | | | | | | | | |
| Nampt | 0.333333 | 0.333333 | 0.75 | | | | | | | | | | | | | | | | | | | | | |
| IR | 1 | 0.333333 | 0.083333 | 0.75 | | | | | | | | | | | | | | | | | | | | |
| pAKT/AKT | 1 | 0.333333 | 0.083333 | 0.75 | 0.083333 | | | | | | | | | | | | | | | | | | | |
| PGC-1a | 0.333333 | 0.333333 | 0.75 | 0.083333 | 0.75 | 0.75 | | | | | | | | | | | | | | | | | | |
| FOXO | 0.333333 | 1 | 0.416667 | 0.75 | 0.416667 | 0.416667 | 0.75 | | | | | | | | | | | | | | | | | |
| pAMPK/AMP | 0.333333 | 1 | 0.416667 | 0.75 | 0.416667 | 0.416667 | 0.75 | 0.083333 | | | | | | | | | | | | | | | | |
| OXPHOS Complex I | 0.75 | 0.75 | 0.333333 | 0.916667 | 0.333333 | 0.333333 | 0.916667 | 0.333333 | 0.333333 | | | | | | | | | | | | | | | |
| OXPHOS Complex II | 0.75 | 0.75 | 0.333333 | 0.916667 | 0.333333 | 0.333333 | 0.916667 | 0.333333 | 0.333333 | 0.416667 | | | | | | | | | | | | | | |
| OXPHOS Complex III | 0.916667 | 0.916667 | 0.75 | 0.75 | 0.75 | 0.75 | 0.75 | 0.75 | 0.75 | 0.333333 | 1 | | | | | | | | | | | | | |
| OXPHOS Complex IV | 0.916667 | 0.916667 | 0.75 | 0.75 | 0.75 | 0.75 | 0.75 | 0.75 | 0.75 | 0.333333 | 1 | 0.083333 | | | | | | | | | | | | |
| PC | 0.75 | 0.75 | 0.333333 | 0.916667 | 0.333333 | 0.333333 | 0.916667 | 0.333333 | 0.333333 | 0.083333 | 0.416667 | 0.333333 | 0.333333 | | | | | | | | | | | |
| PA | 0.333333 | 1 | 0.416667 | 0.75 | 0.416667 | 0.416667 | 0.75 | 0.083333 | 0.083333 | 0.333333 | 0.333333 | 0.75 | 0.75 | 0.233333 | | | | | | | | | | |
| Colesterol | 1 | 0.333333 | 0.083333 | 0.75 | 0.083333 | 0.083333 | 0.75 | 0.416667 | 0.416667 | 0.333333 | 0.333333 | 0.75 | 0.75 | 0.35 | 0.35 | | | | | | | | | |
| HDL | 0.916667 | 0.916667 | 0.75 | 0.75 | 0.75 | 0.75 | 0.75 | 0.75 | 0.75 | 1 | 0.333333 | 0.416667 | 0.416667 | 0.783333 | 0.683333 | 0.45 | | | | | | | | |
| LDL | 0.333333 | 1 | 0.416667 | 0.75 | 0.416667 | 0.416667 | 0.75 | 0.083333 | 0.083333 | 0.333333 | 0.333333 | 0.75 | 0.75 | 0.35 | 0.083333 | 0.133333 | 0.516667 | | | | | | | |
| VLDL | 1 | 0.166667 | 0.166667 | 0.5 | 0.166667 | 0.166667 | 0.5 | 1 | 1 | 0.5 | 0.5 | 1 | 1 | 0.766667 | 0.9 | 0.3 | 0.366667 | 0.666667 | | | | | | |
| Triglicérides | 0.416667 | 0.083333 | 0.333333 | 0.333333 | 0.333333 | 0.333333 | 0.333333 | 1 | 1 | 0.75 | 0.75 | 0.916667 | 0.916667 | 0.9 | 0.833333 | 0.266667 | 0.433333 | 0.9 | 0.2 | | | | | |
| Insulina | 0.75 | 0.75 | 0.916667 | 1 | 0.916667 | 0.916667 | 1 | 0.916667 | 0.916667 | 0.75 | 0.75 | 0.333333 | 0.333333 | 0.683333 | 0.683333 | 0.95 | 0.233333 | 0.95 | 0.833333 | 0.766667 | | | | |
| Glicemia | 0.75 | 0.75 | 0.333333 | 0.916667 | 0.333333 | 0.333333 | 0.916667 | 0.333333 | 0.333333 | 0.416667 | 0.083333 | 1 | 1 | 0.35 | 0.083333 | 0.233333 | 0.35 | 0.133333 | 1 | 0.833333 | 0.45 | | | |
| KITT | 0.166667 | 0.333333 | 0.833333 | 0.166667 | 0.833333 | 0.833333 | 0.166667 | 0.5 | 0.5 | 0.666667 | 1 | 0.666667 | 0.666667 | 0.266667 | 0.266667 | 1 | 0.5 | 0.633333 | 0.2 | 0.25 | 1 | 0.5 | | |
| GTT | 0.916667 | 0.916667 | 0.75 | 0.75 | 0.75 | 0.75 | 0.75 | 0.75 | 0.75 | 1 | 0.333333 | 0.416667 | 0.416667 | 0.95 | 0.516667 | 0.683333 | 0.083333 | 0.683333 | 0.833333 | 0.766667 | 0.083333 | 0.233333 | 0.9 | |
| IDE | 0.083333 | 0.416667 | 1 | 0.333333 | 1 | 1 | 0.333333 | 0.333333 | 0.333333 | 0.75 | 0.75 | 0.916667 | 0.916667 | 0.45 | 0.083333 | 0.783333 | 0.95 | 0.233333 | 0.5 | 0.366667 | 0.516667 | 0.233333 | 0.133333 | 0.683333 |
